# A method to improve the reproducibility of findings from epigenome- and transcriptome-wide association studies

**DOI:** 10.1101/2023.03.29.534761

**Authors:** Edwin JCG van den Oord, Jerry D Guintivano, Karolina A. Aberg

## Abstract

Reproducibility is a cornerstone of scientific progress. In epigenome- and transcriptome-wide association studies (E/TWAS) failure to reproduce may be the result of false discoveries. Whereas multiple methods exist to control false discoveries due to sampling error, minimizing false discoveries due to outliers and other data artefacts remains challenging. We propose a robust E/TWAS approach that outperforms alternative methods to improve reproducibility such as split-half replication. Furthermore, robust E/TWAS results in only a minor loss of power if there are no outliers and can in the presence of outliers, likely a more realistic scenario, even be more powerful than regular E/TWAS.

## BRIEF REPORT

Reproducibility is a cornerstone of scientific progress[1]. In the context of high dimensional biological investigations such as epigenome- and transcriptome-wide association studies (E/TWAS), failure to reproduce may be the result of a lack of generalizability of findings to other study populations, false discoveries due to sampling fluctuations, and false discoveries due to outliers or other data artefacts. To generalize findings, studies need to be performed other populations. False discoveries due to sampling fluctuations can effectively be controlled using standard multiple testing corrections. It is less clear how to best eliminate false discoveries due to outliers and other data artefacts. Replicating findings in independent samples is an option. However, these samples may not always be available, and even if they are, for statistical and pragmatic reasons it will be better to avoid replicating false discoveries as much as possible. The availability of efficient methods to eliminate false discoveries due data artefacts may be particularly important for E/TWAS where many markers are tested for association. That is, although results may be sound for the vast majority of tested biological markers, such false discoveries may be disproportionally over-represented among the top findings as data artefacts may increase the chance of artificially small P values.

In this article, we propose a method for eliminating false discoveries due to outliers and other data artefacts that we call robust E/TWAS. This method involves (i) partitioning the sample in *k* equal and non-overlapping folds, (ii) perform separate association studies in each fold, (iii) perform a signed meta-analysis across all *k* folds using Stouffer’s Z-score method[2] after transforming T statistics from each fold into Z statistics, and (iv) declare significance after controlling for multiple testing using the meta-analysis P values as input.

We performed simulations to evaluate the robust E/TWAS method. We studied either univariate or bivariate outliers. Univariate outliers affect both the outcome or the biological marker (e.g., the methylation site or transcript) but not both variables in the same individual. Bivariate outliers involve outlying values for both the outcome and the biological marker in the same individual. We perform 10,000 simulations for a marker with a sample size of 250 and assuming that the outcome and marker are not associated. We either assume five univariate outliers or one bivariate outlier (bivariate outliers will be rarer than univariate outliers). Significance tests are performed allowing for a Type I error of α=0.05, meaning that the null hypothesis should be rejected in 5% of the simulations if false discoveries are controlled properly. For the sake of comparison, we also study the impact of these outliers on regular association testing that analyzes the entire sample at once, and “split-half replication” where the sample is randomly split in a discovery and replication part. For split-half replication, in simulations where the P value is smaller than 0.05 in the discovery sample we perform a “replication” using a one-sided test in the replication sample assuming the same direction of effect as in the discovery sample.

**Figure 1a** shows the results. Without outliers all methods accurately control the Type I error at 0.05 (results not shown). With univariate outliers, Type I errors are slightly lower than the desired 0.05. The exception is the robust E/TWAS with *k*≥ 10 that accurately controls the Type I errors at the 0.05 level. The bivariate outlier appears to have a much bigger impact. The split-half replication method is most sensitive to this outlier. Thus, the bivariate outlier will be present in either the discovery sample or the replication sample. If present in the discovery sample, it will have an increased probability (> 0.05) of being tested in the replication sample where it is expected to “replicate” in 5% of the simulations (α=0.05). If not present in the discovery sample, it will be tested in 5% of the replication samples (α=0.05) where it will have an increased probability (> 0.05) to “replicate” due to the outlier being in the replication sample. Figure 1a shows that to mitigate the effect of a bivariate outlier, rather than split-half replication it is actually better to analyze the entire sample at once as the vast majority of individuals that have no outliers will dilute the effect of the bivariate outlier. However, by far the best results are obtained with robust E/TWAS where the risk of false discoveries decreases when the number of folds increases. As the effect of the bivariate outlier is limited to one fold, robust E/TWAS dilutes its impact as it combines the association evidence from the affected fold with those from all other folds that are not affected by the outlier.

**Figure 1.**
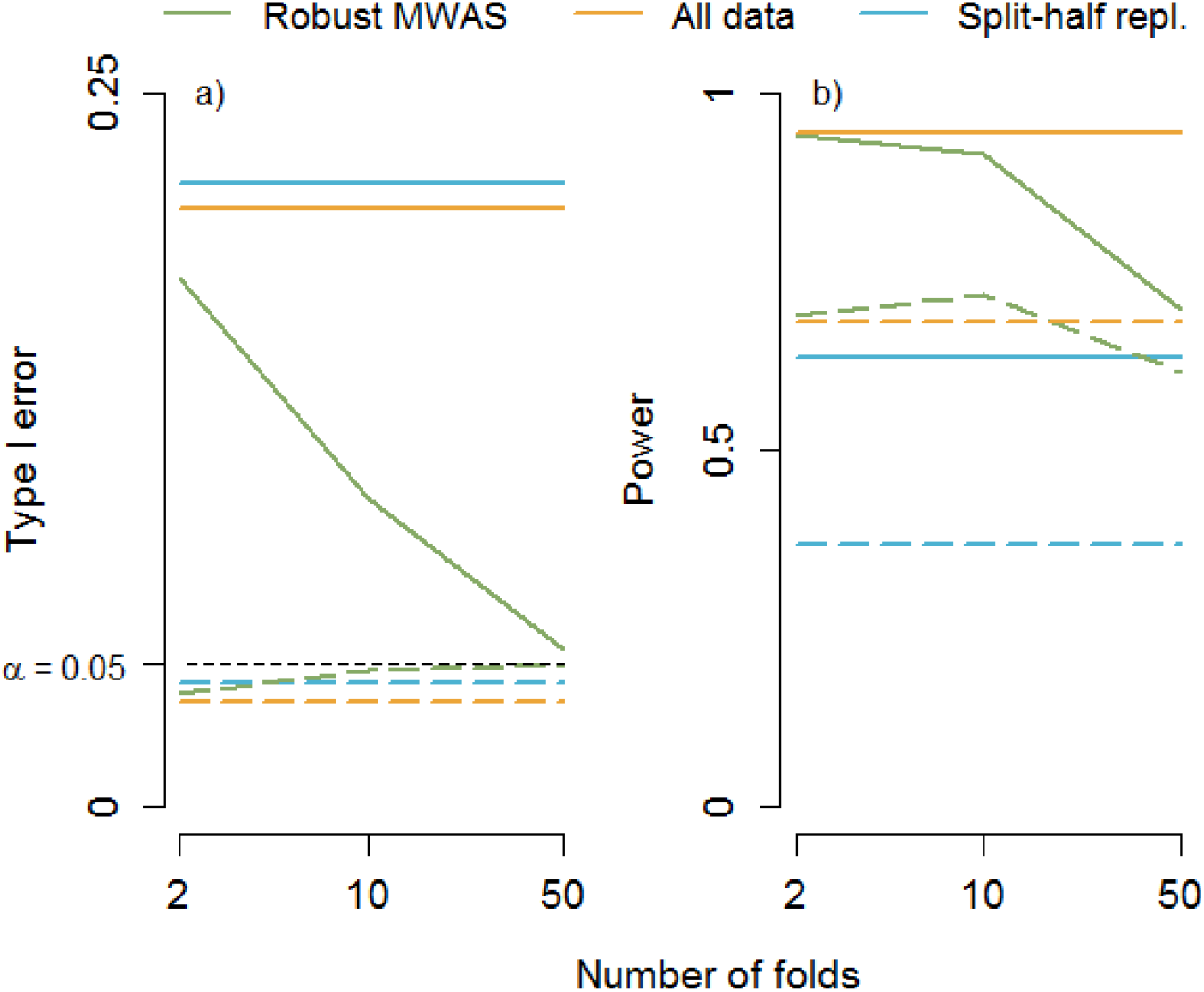
Method comparisons. a) Type I error at α=0.05 (top lines bivariate outliers and dashed bottom lines univariate outliers), b) Power at α=0.001 (top lines no outliers and dashed bottom lines univariate outliers)

The risk of false discoveries needs to be balanced against the risk of false non-discoveries (Type II errors), which is determined by the statistical power. To study power, we repeated the simulations but now assuming a correlation of 0.3 between the outcome and biological marker and use α=0.001 as the threshold for declaring significance. We only studied power in the absence of outliers and in the presence of univariate outliers as these are the common scenarios. Having a bivariate outlier for a marker with a true effect will be rare where, depending on whether the outlier is in line with the association trend, the outlier can either make it more or less likely that the association will be detected. When there are no outliers, robust E/TWAS results in only a slight loss of power compared to analyzing all data at once (**Figure 1b**). In addition, in the presence of univariate outliers robust E/TWAS can even have better power compared to analyzing all data at once. This is because univariate outliers attenuate the association signal. In robust E/TWAS this attenuation is limited to only the folds with the outliers where the unaffected folds mitigate the overall loss in power. The split-half replication has the poorest power. This is because splitting the sample in two parts results in low power in the discovery stage leading to a relatively larger number of simulations where the marker will not proceed to the replication stage and therefore remain undetected.

To illustrate the method with real data we reanalyzed HumanMethylation450 array data for 691 individuals (354 cases and 337 controls, GEO data set GSE42861). After performing quality control as described elsewhere[3], we observed 7 significant findings after a Bonferroni correction for multiple testing. Coefficient lambda (ratio of the median of the observed distribution of the test statistic to the expected median) was 1.126 (Figure S1). This suggested that the vast majority of tests P values were accurate and not influenced by outliers or other data artefacts. The 7 significant results were reanalyzed after removing outliers defined as residual scores (distance between the observed value and the value predicted on the basis of the covariates) with a median absolute deviation (MAD) of >3[4]. Three of the seven significant findings were potentially driven by outliers as removing the outlying observations reduced the P value >100 times. The individuals causing the outlying observations varied across the 7 methylation sites, suggesting that this problem cannot be resolved by eliminating specific individuals from all analyses. Next, we performed a robust MWAS. To keep a fold size of at least 100 individuals, we chose *k*=5 folds. The lambda of 1.083 (Figure S2) was comparable to the lambda of the regular MWAS, suggesting again that the majority of results were not driven by outliers. In the robust MWAS 24 sites reached “suggestive” significance (P < 1×10^-5^). Two of the three sites identified as driven by outliers in the regular MWAS were no longer among these 24 robust MWAS, suggesting it successfully eliminated findings potentially caused by outliers. Two of the four sites that were not flagged as outlier-driven in the regular MWAS were still among the top 24 robust MWAS findings, suggesting it retained part of the sites that were not caused by outliers. The fact that not all four findings that were retained in the regular MWAS were among the top 24 robust MWAS findings may mean that (i) the MAD outlier detection algorithm is not perfect or (ii) because of lower power in the robust MWAS.

In this real data example, a variety of covariates were regressed out. Part of the reason for reduced power in the robust MWAS power may therefore be the loss of the degrees of freedom caused by regressing out covariates in each fold separately. We therefore tested a variant of the robust method that first regressed out effects of covariates in the entire sample, and then performed association testing in each fold separately with residualized outcome scores while charging the loss of degrees of freedom equally across the folds. However, this approximation resulted in deflated P values. We also tried a log data transformation to handle outliers as that will also avoid a loss of degrees of freedom. However, consistent with previous observations that results for transformed data are often not relevant for the original nontransformed data[5], we no longer observed any overlapping results with the original MWAS.

In summary, robust E/TWAS provides an efficient method to improve the reproducibility of findings from high dimensional biological association studies by eliminating false discoveries due to outliers or other data artefacts. It results in only a minor loss of power if there are no outliers but can in the presence of outliers, likely a more realistic scenario, even be more powerful than regular E/TWAS. As it turns a single sample into multiple subsamples, the negative effect of outliers on false discoveries and power is essentially diluted when the association evidence from the affected subsamples is combined with the evidence from the subsamples that are not affected by the outliers. Robust E/TWAS clearly outperformed split-half replication in terms of false discoveries and power. It also provides a more systematic and statistically motivated method for handling outliers compared to traditional methods for outlier detection that would need to be performed post-hoc on a per site basis and involve multiple arbitrary choices. Finally, robust E/TWAS is easy to implement and can be used with any type of association test.

## Supplementary information

Supplementary information accompanies this paper at:

Additional file 1:

Figure S1. QQ-plot MWAS

Figure S2. QQ-plot robust MWAS

## Acknowledgements

Not applicable

## Authors’ contributions

EvdO and KA conceived and designed the experiment. EvdO and JG performed the analyses. EvdO wrote the manuscript. All authors read and approved the final manuscript.

## Availability of data and materials

The scripts used to run the analyses are available from GitHub: https://github.com/ejvandenoord/robust-association. The real data used for illustration is available from GEO under accession number GSE42861.

## Ethics approval and consent to participate

Not applicable

## Consent for publication

Not applicable

