## Supplemental figures for "A method to improve the reproducibility of findings from epigenome- and transcriptome-wide association studies"

Supplemental figures for the article: **A method to improve the reproducibility of findings from epigenome- and transcriptome-wide association studies.**

Figure S1. QQ plot regular MWAS

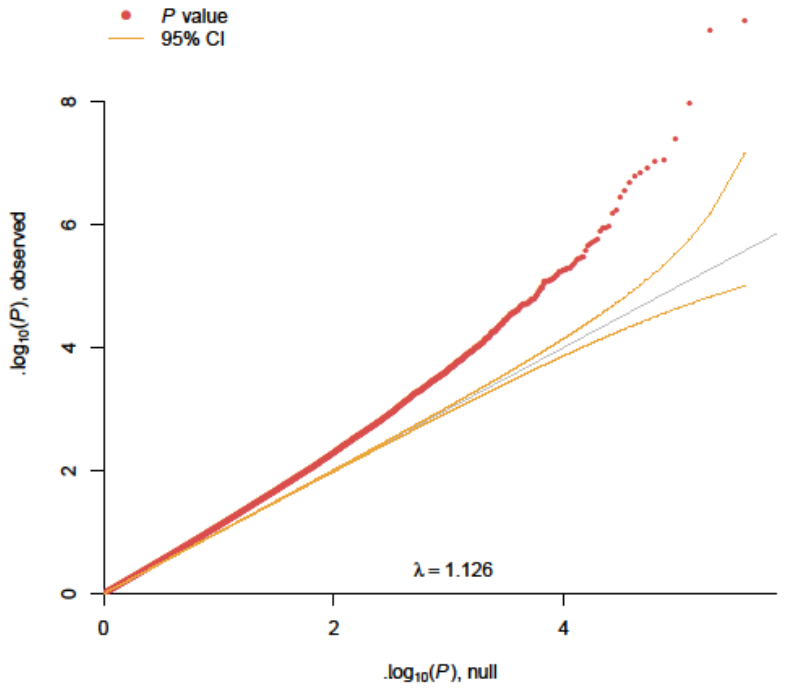

Figure S2. QQ plot for robust MWAS

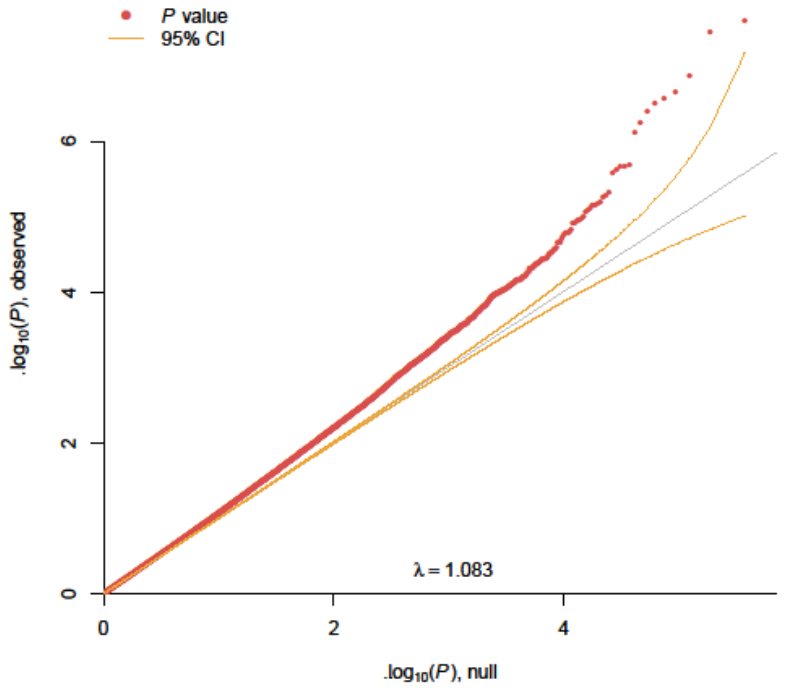
